## Supplementary Figures for "The finger 2 tip loop of Activin A is required for the formation of its non-signaling complex with ACVR1 and type II Bone Morphogenetic Protein receptors"

### Supplementary Materials

A

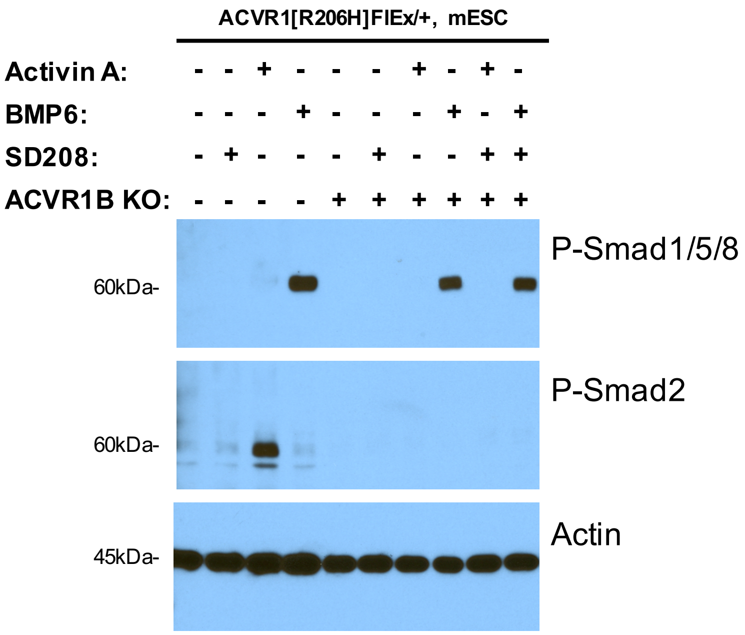


B

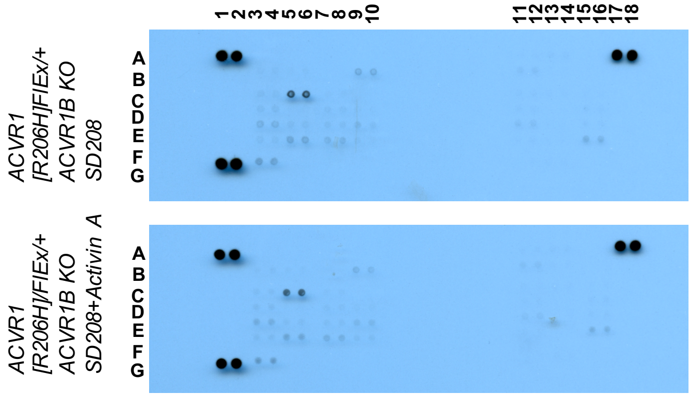

C

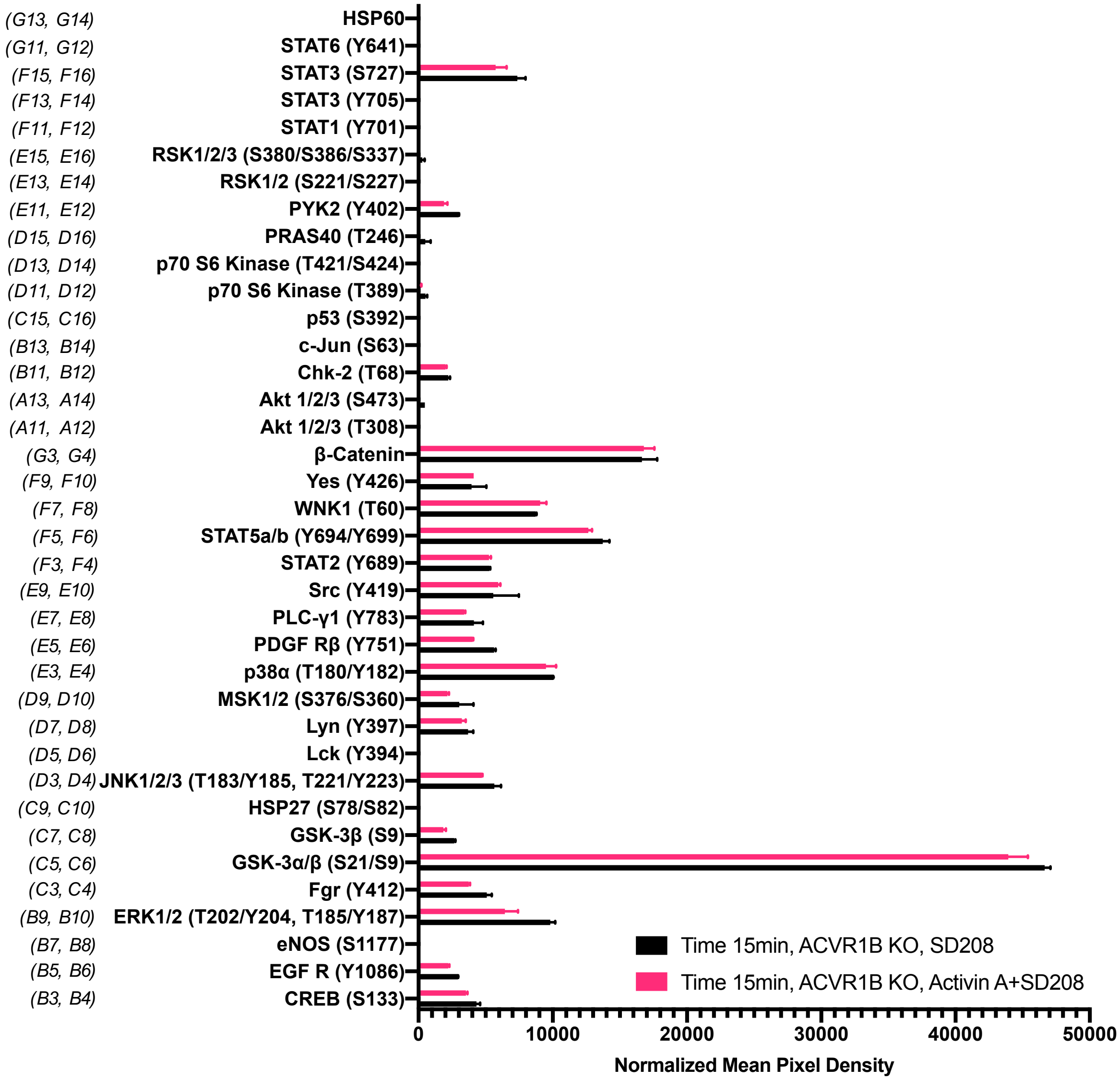


**Supplemental figure 1: The Activin A•ACVR1•type II receptor complex does not transduce signal in mES cells**

(A) Smad-mediated signaling of mouse embryonic stem cells (mES) cells where ACVR1 is not overexpressed was analyzed by immunoblotting using p-Smad1/5/8 and p-Smad2/3 antibodies. In order to isolate signaling induced by Activin A only to the complex that it can form with ACVR1, mES *Acvr1b* knockout (KO) cells were generated and used in comparison to the regular mES cells. mES cells were starved for 3 hours in the presence of 20nM SD208. Then, signaling was induced with 10nM Activin A or 10nM BMP6 in the presence of 20nM SD208 for 15min. Both *Acvr1b* KO and SD208 inhibit Activin A-induced Smad2/3 phosphorylation but not BMP6-induced Smad1/5/8 phosphorylation. (B) Membrane-based sandwich immunoassay analysis of kinase phosphorylation (RnD Systems Proteome Profiler Human Phosphokinase Array Kit) was applied to the same cellular lysates utilized on panel A. (D) Quantitative analysis of Human Phospho-Kinase Array blots shown in panel B. The Activin A•ACVR1•type II receptor complex formed in mES *Acvr1b* KO cells didn't not directly induce downstream phosphorylation of the kinases included in this panel indicating that the Activin A•ACVR1•type II receptor complex doesn't transduce signal.

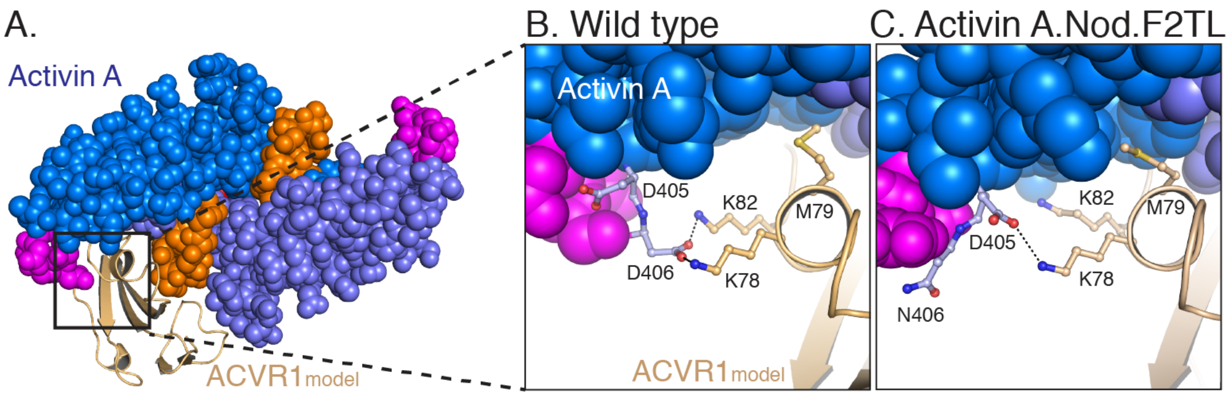


**Supplemental figure 2: Model of Activin A:ACVR1 structure**

(A) Activin A, from its structure with Follistatin 288 (2BOU) (Thompson, Lerch, Cook, Woodruff, & Jardetzky, 2005), was aligned into GDF11 structure from its ternary complex with TGFBR1 and ACVR2B (6MAC) (Goebel et al., 2019). The ACVR1 model was aligned to TGFBR1 in the TGFBR1:GDF11:Acvr2B complex to give the energy minimized model. (B) Closer examination of the F2TL interaction in this model clearly shows Activin A F2TL residue D406 interacting electrostatically with both ACVR1 residues K78 and K82. (C) Substitution of Nodal F2TL into Activin A shows that F2TL coordination is disrupted.

Goebel, E. J., Corpina, R. A., Hinck, C. S., Czepnik, M., Castonguay, R., Grenha, R., . . . Thompson, T. B. (2019). Structural characterization of an activin class ternary receptor complex reveals a third paradigm for receptor specificity. *Proc Natl Acad Sci U S A, 116*(31), 15505-15513. doi:10.1073/pnas.1906253116

Thompson, T. B., Lerch, T. F., Cook, R. W., Woodruff, T. K., & Jardetzky, T. S. (2005). The structure of the follistatin:activin complex reveals antagonism of both type I and type II receptor binding. *Dev Cell, 9*(4), 535-543. doi:10.1016/j.devcel.2005.09.008

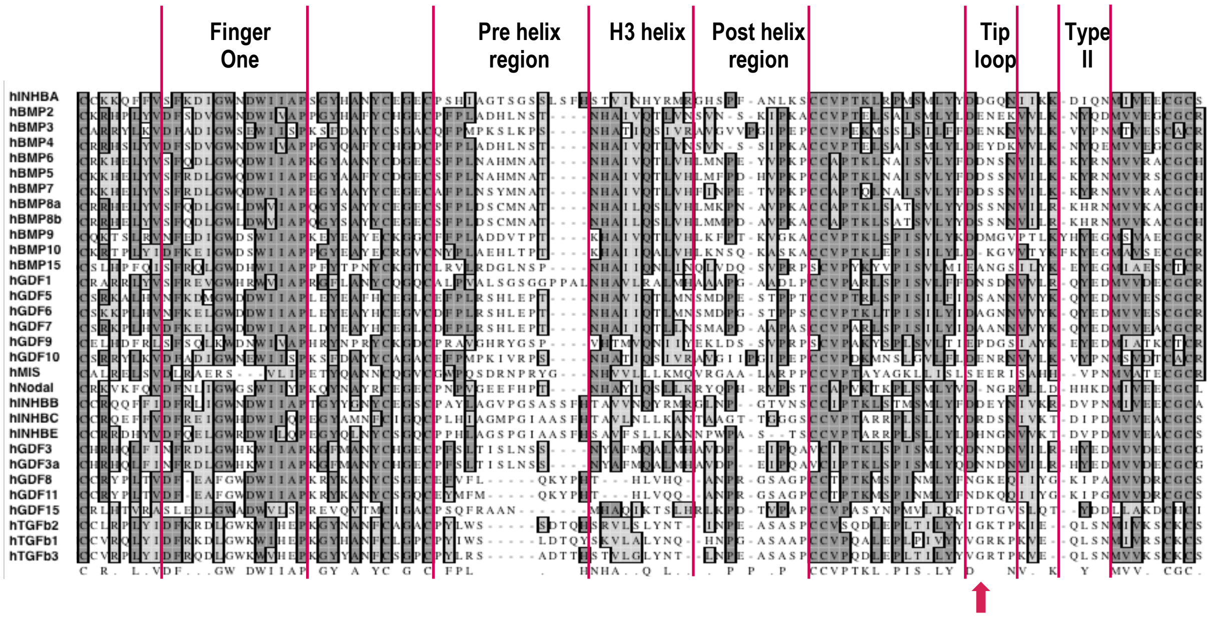


**Supplemental figure 3: Sequence alignment of mature ligands in human TGFβ family**
Sequences of mature human TGFβ family ligands were aligned in MacVector with ClustalW. Structural elements involved in receptor binding are highlighted as follows: finger one, pre-helix region, H3 helix (central helix), post-helix region, tip loop (F2TL), and site of sequence diversity coding for type II receptor binding specificity (type II). In looking for regions of sequence diversity within the family, the pre-helix, post-helix and F2TL regions were chosen for substitutional mutagenesis analysis of Activin A. Position 406 is marked with a red arrow.

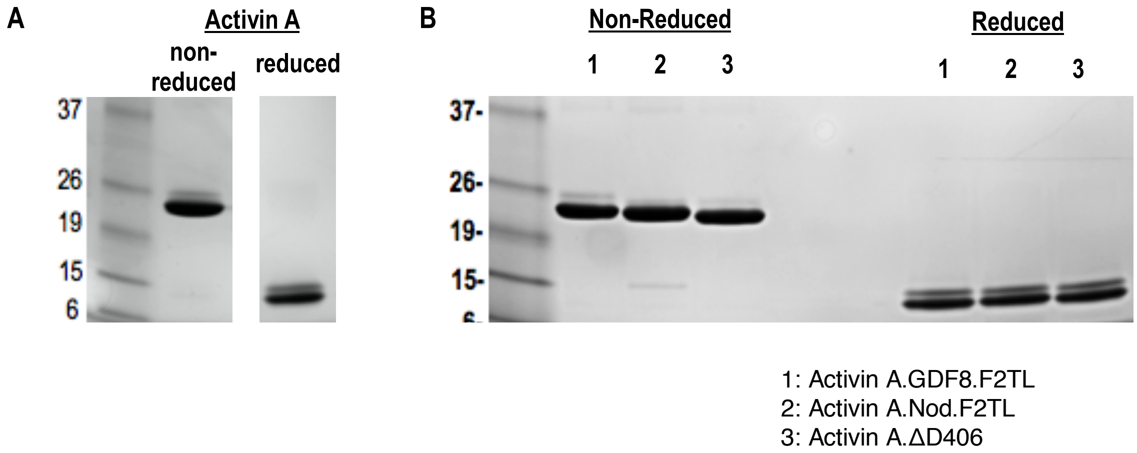


**Supplemental figure 4: Purified Activin and Activin A F2TL muteins**
Activin A was purified from CHO-K1 cell supernatants by heparin affinity and ion exchange chromatography. The mature form of Activin A and Activin A muteins was separated by reverse phase chromatography, and products were analyzed by Coomassie blue stained SDS-PAGE gels. (A) Activin A and (B) Activin A F2TL muteins are shown under reducing and non-reducing conditions.

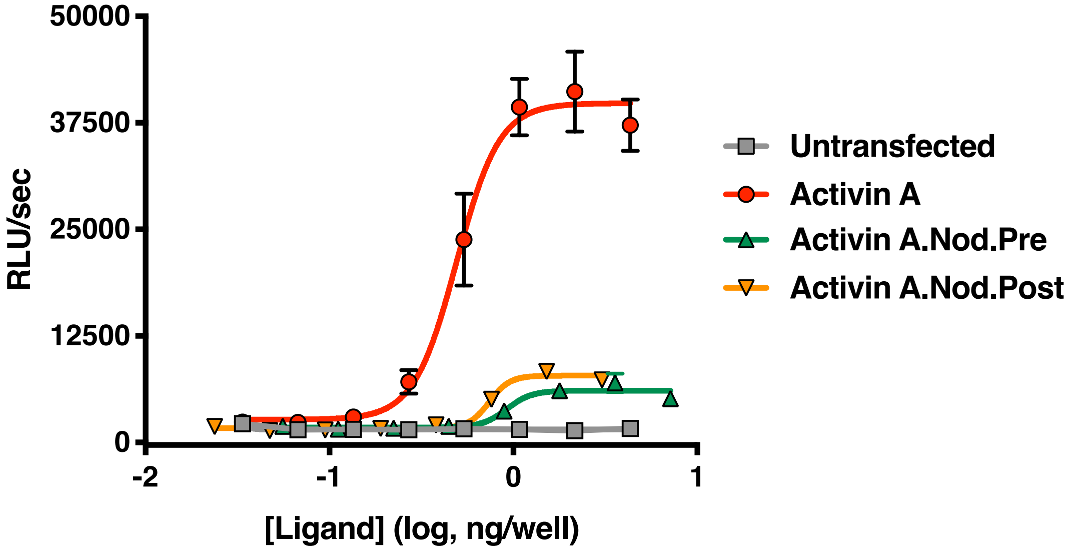


**Supplemental figure 5: Activin A with pre-helix and post-helix substitutions from Nodal display reduced ability to activate the Smad2/3 pathway**

Supernatants from CHO cells expressing Activin A with the pre or post-helix sequence from human Nodal were tested for activity in HEK293 Smad2/3 reporter cells. Both the Activin A.Nod.Pre and Activin A.Nod.Post supernatants display reduced activity compared to Activin A (R&D Systems). The data presented are representative three independent biological and technical replicates.

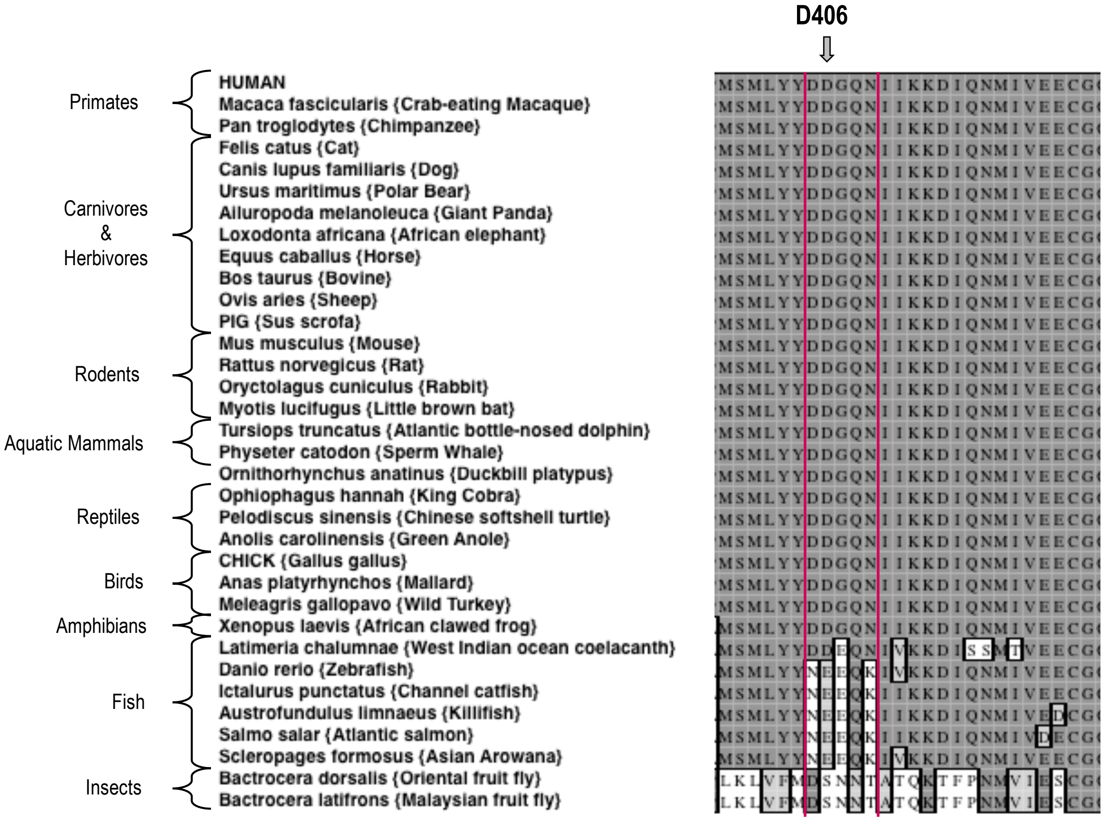


**Supplemental figure 6: Sequence alignment of Activin A from various species**
A Clustal W alignment with the F2TL sequence bracketed in red. Human Activin A aspartic acid at position 406, D406, is evolutionarily well conserved in Activin A from different vertebrates. Divergence of this sequence is apparent in Activin A from the majority of fish species examined here.

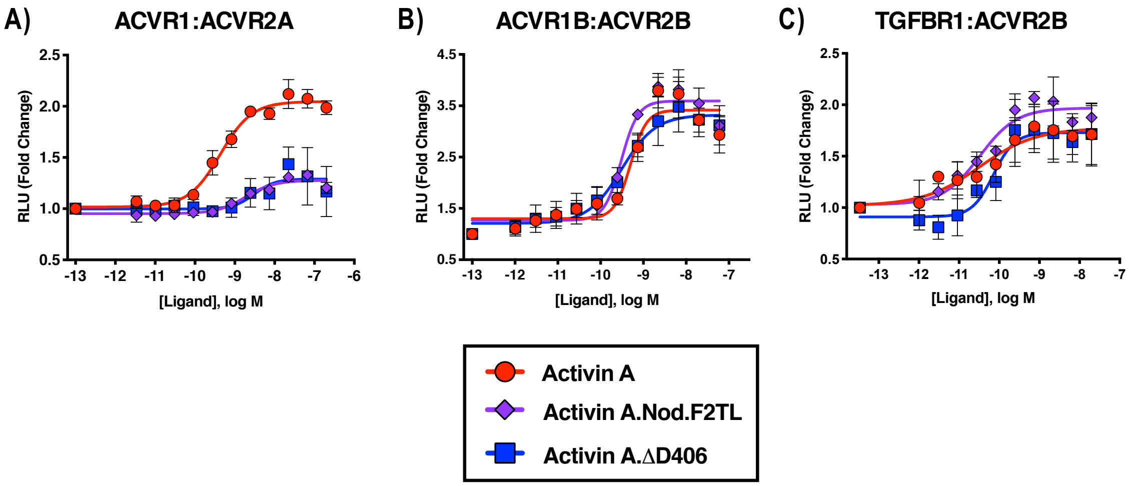


**Supplemental figure 7: Activin A F2TL muteins lose binding to ACVR1**

U20S cells expressing split beta-galactosidase fusions of corresponding type I and type II receptors were treated with a dose response of Activin A ligands. Type I receptor binding was measured by luminescence. (A) The finger two tip loop mutants have reduced ability to dimerize ACVR1 with ACVR2A, while retaining wild type capacity to dimerize (B) ACVR1B with BMPR2 and (C) TGFBR1 with ACVR2B. The data presented are representative of three independent biological and technical replicates.

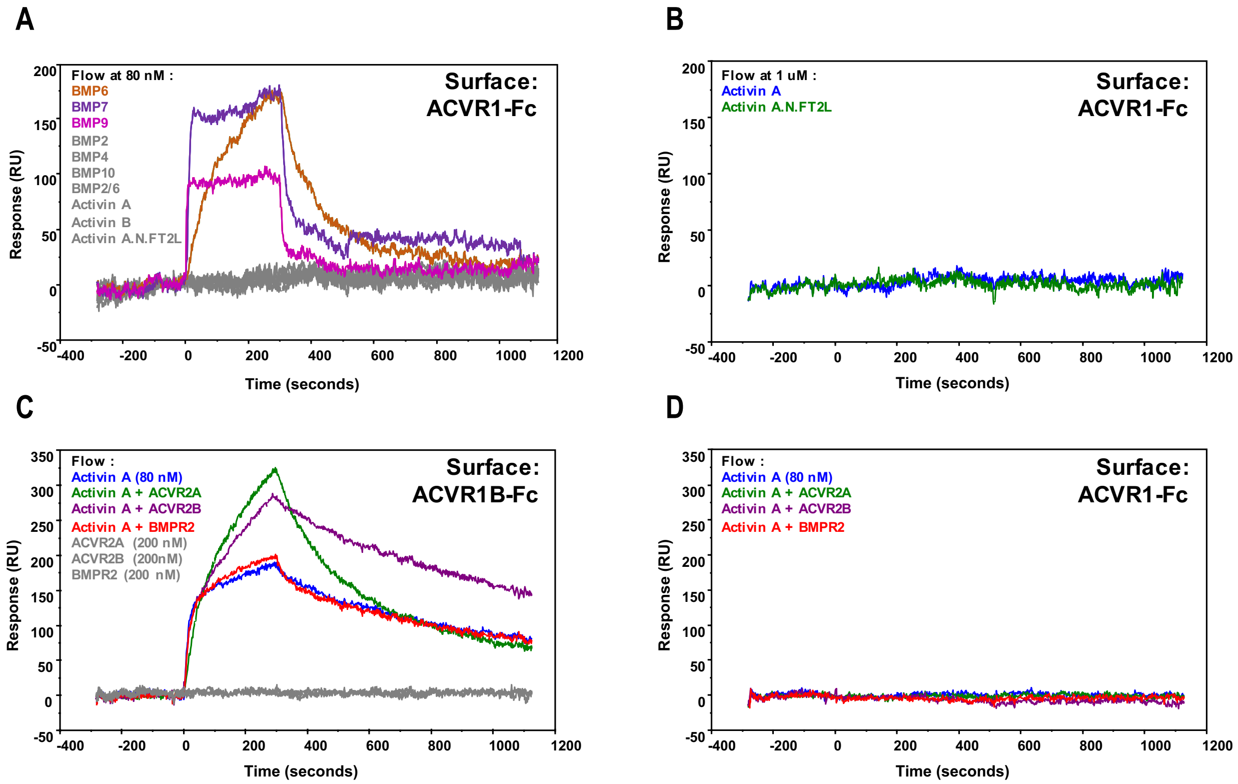


**Supplemental Figure 8: Activin A•ACVR1 complex cannot be detected using SPR**
For (A) and (B), 1000 RU of either ACVR1B-Fc or ACVR1-Fc were captured on a sensor chip. Ligands at 80 nM (A) or 1 μM (B) were injected, including Activin A, Activin B, Activin A.Nod.F2TL, BMP2, BMP4, BMP7, BMP9, BMP10, and BMP2/6 over ACVR1-Fc. ACVR1-Fc binds to BMP6, BMP7, and BMP9, but does not form a complex with Activin A, Activin B, Activin A.Nod.F2TL, BMP2, BMP4, BMP10, and BMP2/6. Ligands and corresponding binding curves are color matched. For (C) and (D), complexes of Ligand•Type II receptor extracellular domain were formed at 1:2.5 ratio and injected over ACVR1B-Fc and ACVR1-Fc. C) Activin A•ACVR1B-Fc complex is detectable by SPR either alone or in the presence of monomeric extracellular domain of ACVR2A or ACVR2B. Activin A•ACVR2A and Activin A•ACVR2B, but not Activin A•BMPR2, complexes bound ACVR1-Fc at a stoichiometric ratio. (D) In contrast, none of the tested Activin A•Type II receptor complexes bound ACVR1-Fc. Ligand +/- Type II receptor and corresponding binding curves are color matched. The data shown here is a representative of two biochemical and two technical replicates.

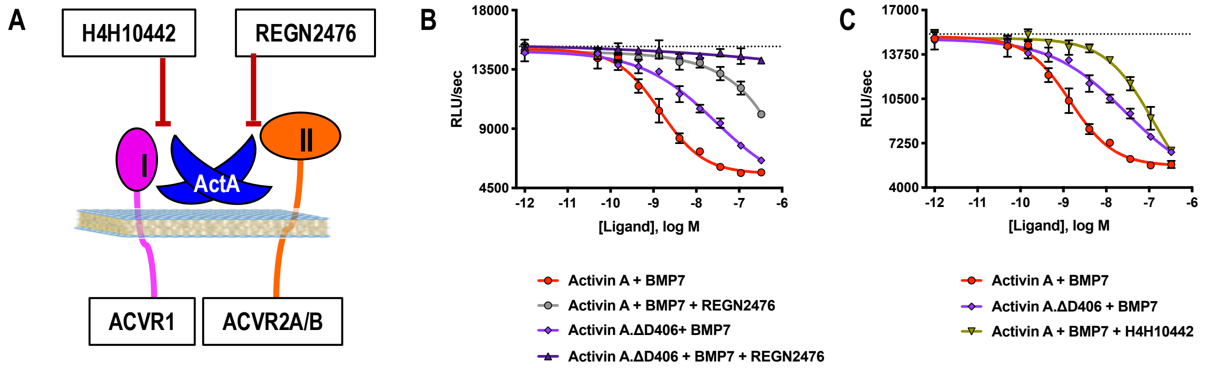


**Supplemental figure 9: Activin A.ΔD406 is 15-fold less effective than Activin A at inhibiting BMP7-induced Smad1/5/8 signaling through ACVR1**

(A) Schematic showing H4H10442 blocking Activin A (ActA) binding to type I receptor and REGN2476 preventing Activin A from binding to the type II receptor. (B, C) Varying concentrations of Activin A or Activin A.ΔD406 were mixed with a constant concentration of BMP7 (12nM) and applied to HEK293 cells stably transfected with the Smad1/5/8 reporter construct driving firefly luciferase. (B) Using REGN2476, we show the remaining inhibition of BMP7 by Activin A.ΔD406 is lost when type II receptor binding is blocked. (C) Inhibition of type I receptor binding of wild type Activin A shows further reduction in BMP inhibition compared to Activin A.ΔD406 alone. Inhibition of BMP7 is reduced ~15-fold with Activin A.ΔD406 compared to ~60-fold with the Activin A:H4H10442 complex. (The $IC_{50}$s of Activin A and Activin A.ΔD406 are $1.4 \times 10^{-9}$ M and $2.0 \times 10^{-8}$ M, respectively.) The data presented are representative of three independent biological and technical replicates.

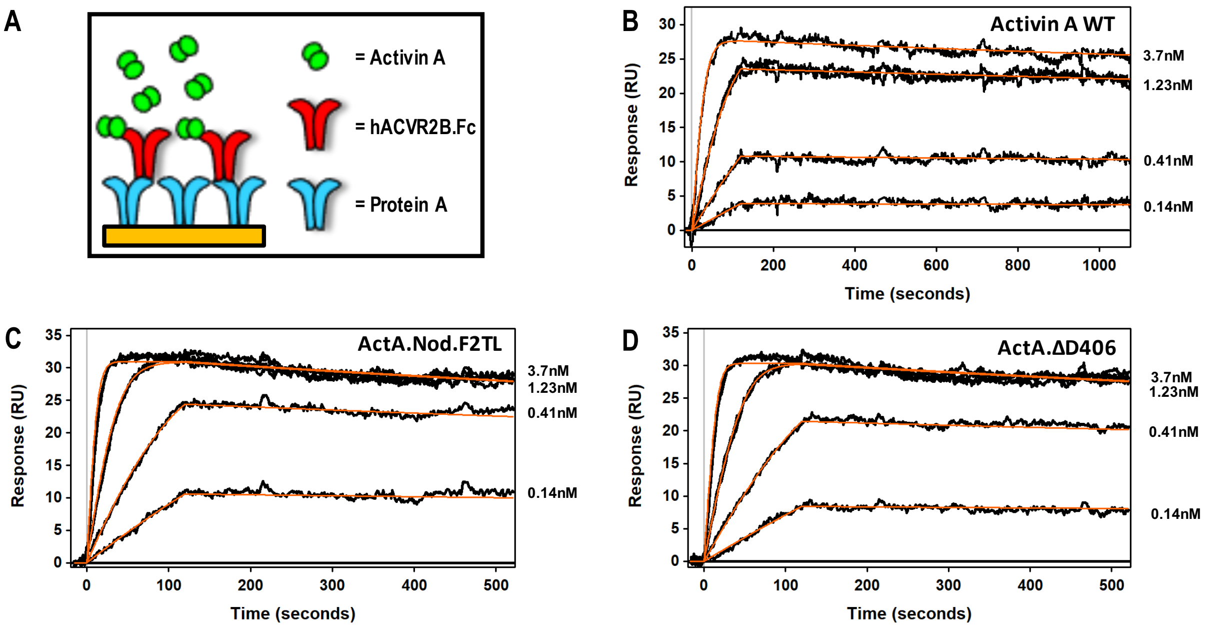


E

| Capture Surface | Dimer Captured (RU) | Reagent Tested | 100nM Reagent Bound (RU) | ka (1/Ms) | kd (1/s) | KD (M) | t½ (min) |
|---|---|---|---|---|---|---|---|
| ACVR2B.Fc | 118 ±1 | ActivinA WT | 30 | 1.89E+07 | 8.46E-05 | 4.49E-12 | 137 |
| | 117 ±0.9 | ActA.Nod.F2TL | 33 | 5.29E+07 | 2.70E-04 | 5.11E-12 | 43 |
| | 116 ±0.8 | ActA.ΔD406 | 33 | 5.50E+07 | 2.54E-04 | 4.62E-12 | 46 |

**Supplemental figure 10: Activin A finger 2 tip loop muteins do not display altered type II receptor binding**

(A) Schematic representation of Surface Plasmon Resonance binding assay. ACVRIIB.Fc was captured on a sensor chip and measured its binding to (B) Activin A WT, (C) ActA.Nod.T2L, (D) ActA.ΔD406. (E) Binding constants of mature Activin A and Activin A mutein proteins to Protein A captured ACVR2B Fc. Mutations to Activin A F2TL do not disturb type II receptor binding site. The data presented are representative of three independent biochemical replicates.

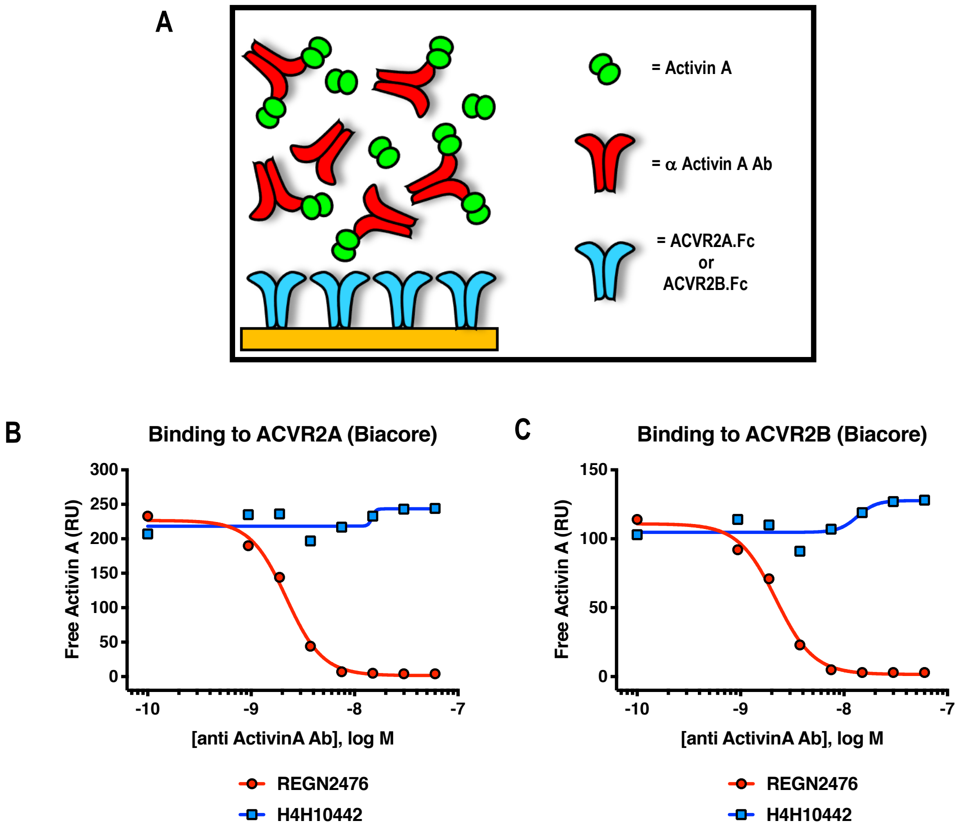


**Supplemental figure 11: Anti-Activin A antibody REGN2476 blocks binding of Activin A to type II receptors ACVR2A and ACVR2B**

(A) Schematic representation of Surface Plasmon Resonance binding assay. SPR was utilized to evaluate binding of Activin A to surface-immobilized ACVR2A-Fc or ACVR2B-Fc, in the presence of increasing amounts of anti-Activin A antibodies. (B, C) Anti-Activin A antibody H4H10442 and REGN2476 at concentration range 0.94 nM-60 nM were pre-mixed with 5nM Activin A and injected over (B) ACVR2A or (C) ACVR2B Fc fusion protein coupled chip surfaces. H4H10442 allows Activin A binding to ACVR2A and ACVR2B surfaces. REGN2476 directly blocks Activin A binding to both type II receptors.

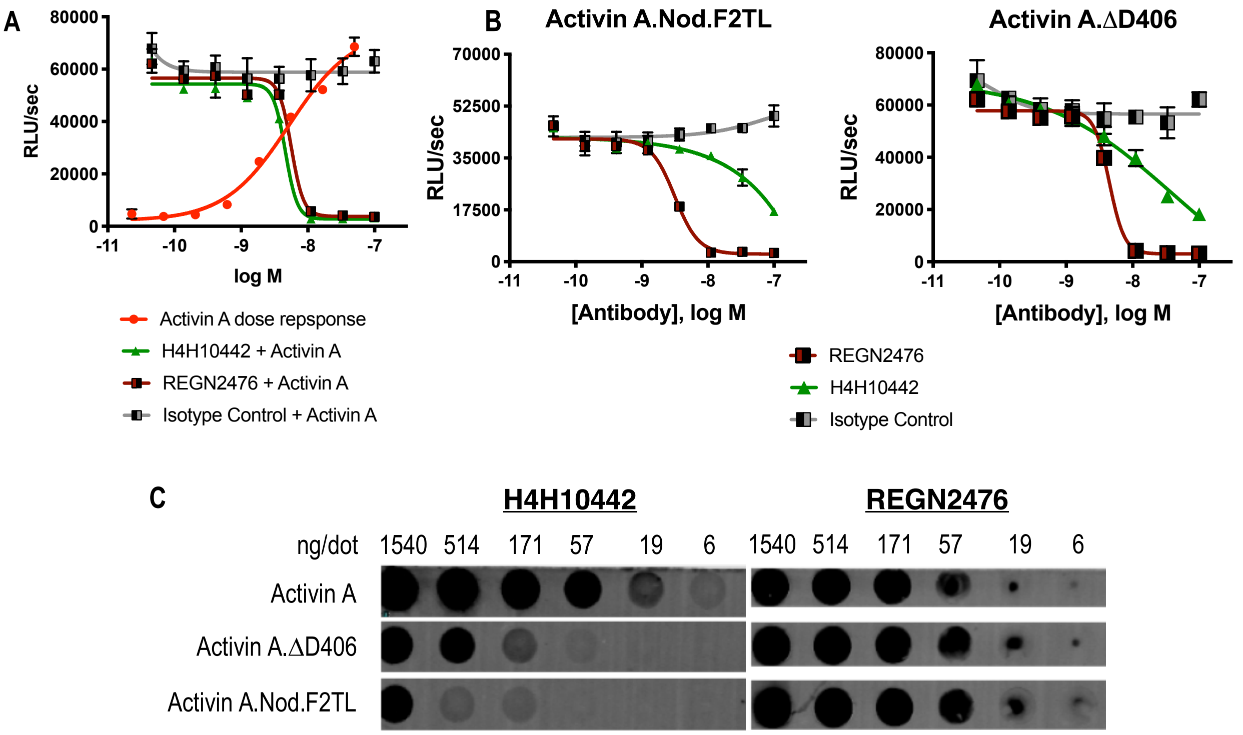


D

| Capture Surface | Dimer Captured (RU) | Reagent Tested | 20nM Activin A bound (RU) | ka (1/Ms) | kd (1/s) | KD (M) | t½ (min) |
|---|---|---|---|---|---|---|---|
| H4H10442 | 181 | Activin A | 41 | 1.44E+07 | 5.00E-05 | <3.48E-12 | >231 |
| REGN2476 | 109 | | 25 | 2.30E+07 | 5.00E-05 | 3.97E-12 | 77 |

**Supplemental figure 12: The anti-Activin A blocking antibody H4H10442 inhibits Activin A by binding to its finger 2 tip loop (F2TL) region**

(A) Both H4H10442 and REGN2476 anti-Activin A antibodies block Activin A (10nM) signaling through the Smad2/3 pathway in HEK293 CAGA-luciferase reporter cells. (B) In HEK293 Smad2/3 luciferase reporter cells, H4H10442 is a less effective inhibitor of the Activin A F2TL muteins (10 nM) when compared to wild type Activin A. The data presented are representative of two independent biological and technical replicates. (C) Anti-Activin A monoclonal antibody H4H10442 appears to recognize an epitope that overlaps with the F2TL region of Activin A, as it does not bind as well to either Activin A.ΔD406 or Activin A.Nod.F2TL. The ability of H4H10442 to bind the latter is particularly hampered. In contrast, REGN2476 bound similarly to all three ligands, consistent with the finding that the type II receptor-interacting regions of Activin A.ΔD406 and Activin A.Nod.F2TL have not been altered. For dot blots, purified Activin A and Activin A muteins were serially diluted and applied to PVDF membranes using suction. Membranes were blocked using Odyssey blocking reagent and the hIGg4 was visualized using an IRDye 680RC conjugated goat anti-human secondary antibody (Li-cor). (D) Binding affinities and kinetic constants for binding purified anti-human Activin A monoclonal antibodies H4H10442 and REGN2476 to Activin A were determined using a real-time surface plasmon resonance biosensor at 37°C. Antibodies were captured on anti-human Fc sensor surfaces. Activin A-antibody association rates were measured by injecting 20nM Activin A over the antibody captured surface. Both H4H10442 and REGN2476 bind Activin A with very high affinity, displaying KDs in low picomolar range.

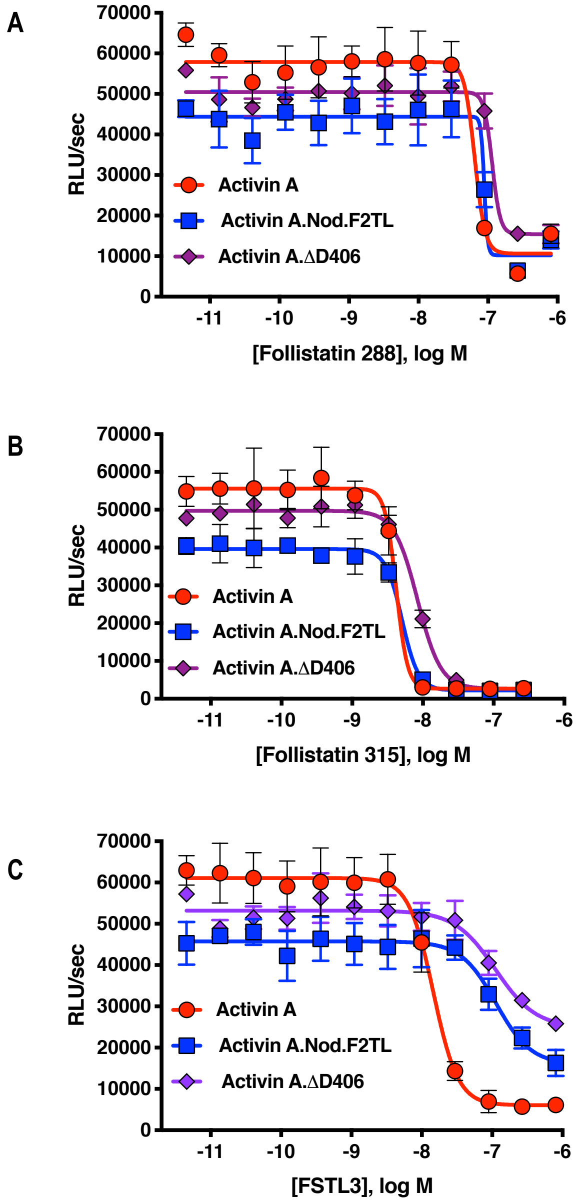


**Supplemental Figure 13: Activin A F2TL muteins are inhibited by Follistatin but show reduced inhibition by FSTL3**

Varying concentrations of Follistatin and FSTL3 were preincubated with a constant concentration (10nM) of Activin A, Activin A.Nod.F2TL or Activin A. Δ.D406. Activity was tested in HEK293 cells harboring a Smad2/3 luciferase reporter. Activity of Activin A, ActivinA.Nod.F2TL and ActivinA. ΔD406 was blocked by both (A) follistatin-288 and (B) follistatin-315. (C) FSTL3 is a less effective inhibitor of ActivinA.Nod.F2TL and ActivinA.ΔD406 when compared to wild type Activin A. The data presented are representative of three independent biological and technical replicates.

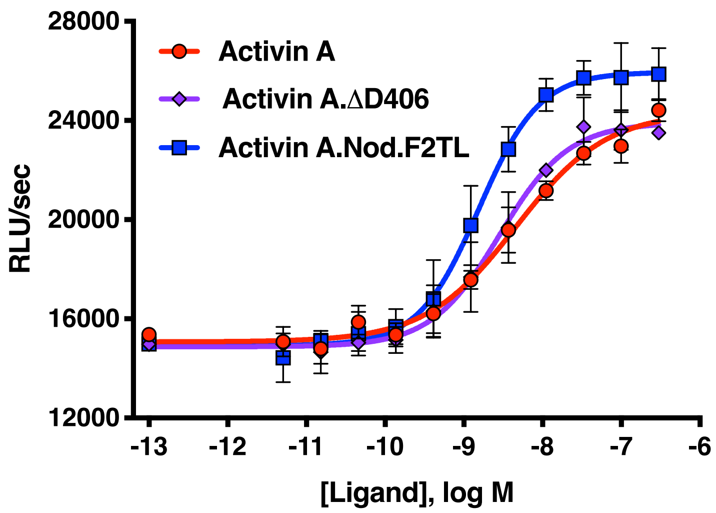


**Supplemental figure 14: Activin A F2TL muteins activate ACVR1[R206H] like wild type Activin A**

HEK293 cells expressing FOP mutant ACVR1 (ACVR1[R206H]) were treated with a dose response of Activin A ligands. Both wild type Activin A and the Activin A F2TL muteins activate Smad1/5/8 pathway via ACVR1[R206H] receptor in the BRE-luciferase reporter assay. Notably, the Activin A.Nod.F2TL mutein is more active than either Activin A or Activin A.ΔD406, consistent with the fact that Activin A.Nod.F2TL is not capable of forming a functional NSC with wild type ACVR1. The data presented are representative of three independent biological and technical replicates.

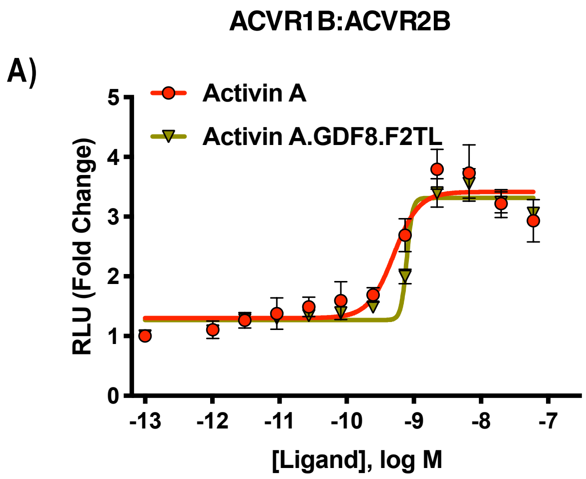


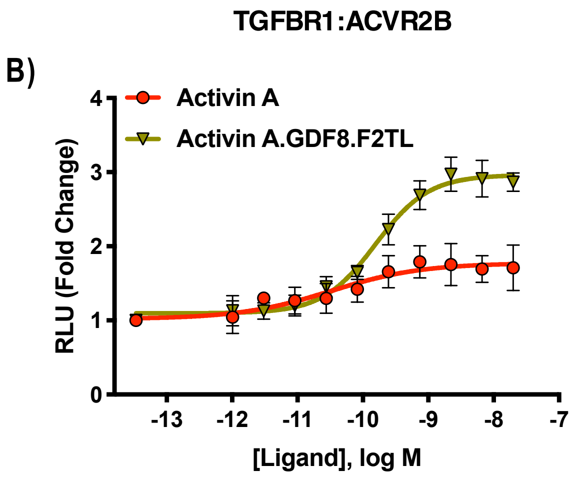


**Supplemental Figure 15: Activin A.GDF8.F2TL mutein binds TGFBR1 better than Activin A**

U20S cells expressing split beta-galactosidase fusions of corresponding type I and type II receptors were treated with a dose response of Activin A.GDF8.F2TL or Activin A. Type 1 receptor binding was measured by luminescence in these receptor dimerization assays. (A) Activin A.GDF8.F2TL retains wild type capacity to dimerize ACVR1B:BMPR2, and (B) Activin A.GDF8.F2TL increases dimerization of the TGFBR1:ACVR2B receptor pairs. The data presented are representative of two independent biological and three technical replicates.

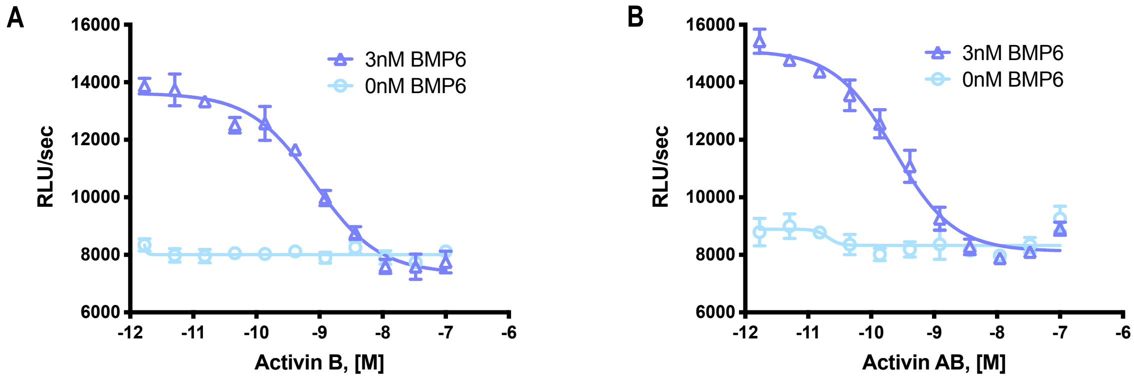


**Supplemental Figure 16: Activin B and Activin AB form a NSC with ACVR1**
HEK293 cells expressing wild type ACVR1 were treated with 3nM BMP6 containing a dose response of either (A) Activin B or (B) Activin AB. Activin antagonism of BMP6 signaling was measured by luminescence from a BRE-luciferase reporter. The data presented are representative of two independent biological and three technical replicates.
